# xCell: Digitally portraying the tissue cellular heterogeneity landscape

**DOI:** 10.1101/114165

**Authors:** Dvir Aran, Zicheng Hu, Atul J. Butte

## Abstract

Tissues are complex milieu consisting of numerous cell-types. Numerous recent methods attempt to enumerate cell subsets from transcriptomes. However, available method used limited source for training and displayed only partial portrayal of the full cellular landscape. Here we present *xCell*, a novel gene-signature based method for inferring 64 immune and stroma cell-types. We harmonized 1,822 pure human cell-types transcriptomes from various sources, employed curve fitting approach for linear comparison of cell-types, and introduced a novel spillover compensation technique for separating between cell-types. Using extensive *in silico* analyses and comparison to cytometry immunophenotyping we show that *xCell* outperforms other methods: http://xCell.ucsf.edu/.

## Introduction

In addition to malignant proliferating cells, tumors are also composed of numerous distinct non-cancerous cell types and activation states of those cell types. This notion, which is termed the tumor microenvironment, has been in the spotlight of research in recent years and is being further explored by novel techniques. The most studied set of non-cancerous cell types are the tumor-infiltrating lymphocytes (TILs). However, these TILs are only part of a variety of innate and adaptive immune cells, stroma cells and many other cell types that are found in the tumor and interact with the malignant cells. This complex and dynamic microenvironment is now recognized to be important both in promoting and inhibiting of tumor growth, invasion, and metastasis [1,2]. Understanding the cellular heterogeneity composing the tumor microenvironment is key for improving existing treatments, the discovery of predictive biomarkers and development of novel therapeutic strategies.

Traditional approaches for dissecting the cellular heterogeneity in liquid tissues are difficult to apply in solid tumors [3]. Therefore, in the last decade, numerous methods have been published for digitally dissecting the tumor microenvironment using gene expression profiles [4–7] (Reviewed in [8]). Recently, multitudes of studies have been published applying published and novel techniques on publicly available resources of tumor samples such as The Cancer Genome Atlas (TCGA) [6, 9–13]. There are two general types of techniques: deconvolving the complete cellular composition, and assessing enrichments of individual cell types.

There are at least seven major concerns that the *in silico* methods could be prone to errors, and cannot reliably portray the cellular heterogeneity of the tumor microenvironment. First, current techniques depend on the expression profiles of purified cell types to identify reference genes and therefore rely heavily on the data source of which the references are inferred from, and could be inclined to overfitting to these data. Second, current methods portray only a very narrow perspective of the tumor microenvironment. The available methods usually focus on a subset of immune cell types, thus not accounting for the further richness of cell types in the microenvironment, including blood vessels and other different forms of cell subsets [14,15]. A third problem is the ability of cancer cells to “imitate” other cell types by expressing immune-specific genes, such as macrophages-like expression pattern in tumors with parainflammation [16]; only a few of the methods take this into account. Fourth, the ability of existing methods to estimate cell abundance have not yet been comprehensively validated in mixed samples. Cytometry is a common method for counting cell types in a mixture, and when performed in combination with gene expression profiling, can allow validation of the estimations. However, in most studies that included cytometry validation, these analyses were performed on only a very limited number of cell types and a limited number of samples [7,13].

A fifth challenge is that deconvolution approaches are prone to many different biases because of the strict dependencies among all cell types that are inferred. This could highly affect reliability in analyzing tumor samples, which are prone to form non-conventional expression profiles. A sixth problem has been raised with inferring an increasing number of closely related cell types [10]. Finally, deconvolution analysis heavily relies on the structure of the reference matrix, which limits its application to the resource used to develop the matrix. One such deconvolution approach is CIBESORT, which is the most comprehensive study to date, allowing the enumeration of 22 immune subsets [7]. Newman et al. performed adequate evaluation across data sources and validated the estimations using cytometry immunophenotyping. However, the shortcomings of deconvolution approaches are apparent in CIBERSORT, which is limited to Affymetrix microarray studies.

On the other hand, gene set enrichment analysis is a simple technique, which can be easily applied across data types and can be quickly applied for cancer studies. Each gene signature is used independently from all other signatures; thus it is protected from the limitations of deconvolution approaches. However, because of this independence, it is many times hard to differentiate between closely related cell types. In addition, gene signature-based methods only provide enrichment scores, and thus do not allow comparison across cell types, and cannot allow insights on the abundance of the cell type in the mixture.

Here, we present *xCell*, a novel method that integrates the advantages of gene set enrichment with deconvolution approaches. We present a compendium of newly generated gene signatures for 64 cell types, spanning multiple adaptive and innate immunity cells, hematopoietic progenitors, epithelial cells and extracellular matrix cells derived from thousands of expression profiles. Using *in silico* mixtures, we transform the enrichment scores to a linear scale, and using a spillover compensation technique we reduce dependencies between closely related cell types. We evaluate these adjusted scores in RNA-seq and microarray data from primary cell types samples from various independent sources. We examine their ability to digitally dissect the tumor microenvironment by *in silico* analyses, and perform the most comprehensive comparison to date with cytometry immunophenotyping. We compare our inferences with available methods and show that scores from *xCell* are more reliable in digital dissection of mixed tissues. Finally, we apply our method on TCGA tumor samples to portray a full tumor microenvironment landscape across thousands of samples. We provide these estimations to the community and hope that this resource will allow researchers gain a better perspective of the complex cellular heterogeneity in tumor tissues.

## Results

### Generating a gene signature compendium of cell-types

To generate our compendium of gene signatures for cell types, we collected gene expression profiles from six sources: the FANTOM5 project, from which we annotated 719 samples from 39 cell types analyzed by the Cap Analysis Gene Expression (CAGE) technique [17]; the ENCODE project, from which we annotated 115 samples from 17 cell types analyzed by RNA-seq [18]; the Blueprint project, from which we annotated 144 samples from 28 cell types analyzed by RNA-seq (http://www.blueprint-epigenome.eu/); the IRIS project, from which we annotated 95 samples from 13 cell types analyzed by Affymetrix microarrays [19]; the Novershtern et al. study, from which we annotated 180 samples from 24 cell types analyzed by Affymetrix microarrays [20]; and the Human Primary Cells Atlas (HPCA), a collection of Affymetrix microarrays composed of many different Gene Expression Omnibus (GEO) datasets, from which we annotated 569 samples from 41 cell types [21] (Figure 1A). Altogether we collected and curated gene expression profiles from 1,822 samples of pure cell types, annotated to 64 distinct cell types and cell subsets (Figure 1B and Supplementary Table 1). Of those, 54 cell types were found in at least 2 of these data sources. For cell types with 5 or more samples in a data source, we left one sample out for testing. All together, 97 samples were left out, and all of the model trainings described below were performed on the remaining 1,725 samples.

**Figure 1.**
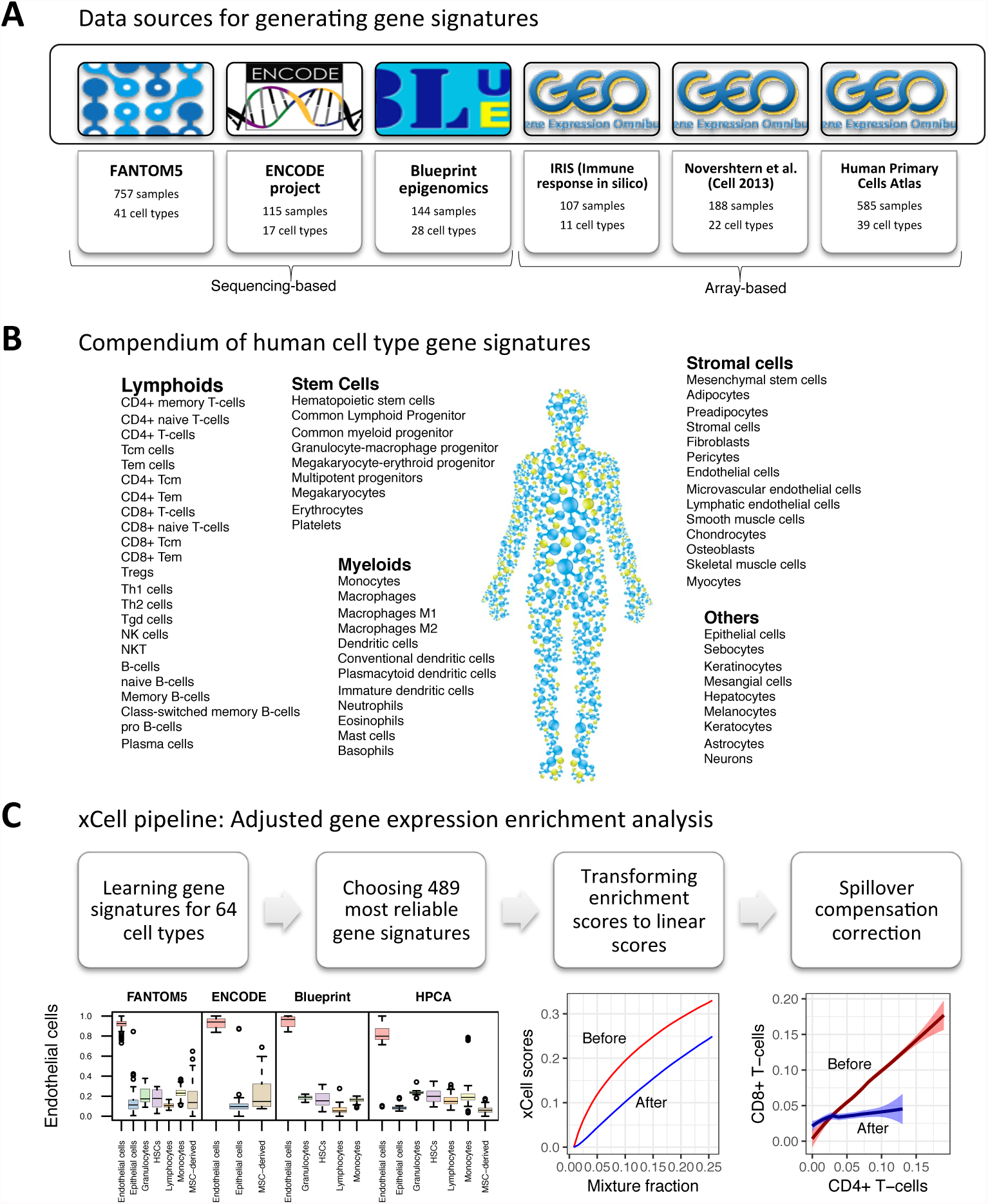
xCell study design. **A.** A summary of Data sources used in the study to generate the gene signatures, showing the number of pure cell types and number of samples curated from them. **B.** Our compendium of 64 human cell types gene signatures grouped into 5 cell types families. **C.** The xCell pipeline: Using the data source and based on different thresholds we learn gene signatures for 64 cell types. Of this collection of 6,573 signatures we choose 489 most reliable cell types, 3 for each cell types from each data source were it is available. The raw score is then the average ssGSEA score of all signatures corresponding to the cell type. Using simulations of gene expression for each cell type we learn a function to transform the non-linear association between the scores to a linear scale. Using the simulations we also learn the dependencies between cell types scores and apply a spillover compensation method to adjust the scores.

Our strategy for selecting reliable cell type gene signatures is shown in Figure 1C and Supplementary Figure 1 (see Methods for full description and technical details). For each data source independently we identified genes that are overexpressed in one cell type compared to all other cell types. We applied different thresholds for choosing set of genes to represent the cell type gene signatures; hence from each source, we generated dozens of signatures per cell type. This scheme yielded 6,573 gene signatures corresponding to 64 cell types. Importantly, since our primary aim is to develop a tool for studying the cellular heterogeneity in the tumor microenvironment, we applied a methodology we previously developed [16] to filter out genes that tend to be overexpressed in a set of 634 carcinoma cell lines from the Cancer Cell Line Encyclopedia (CCLE) [22].

Next, we used single-sample gene set enrichment analysis (ssGSEA) to score each sample based on all signatures. ssGSEA is a well-known method for aggregating a single score of the enrichment of a set of genes in the top of a ranked gene expression profile [23]. To choose the most reliable signatures we tested their performance in identifying the corresponding cell type in each of the data sources. To prevent overfitting, each signature learned from one data source was tested in other sources, but not in the data source it was originally inferred. To reduce biases resulting from a small number of genes and from the analysis of different platforms, instead of one signature per cell type, the top three ranked signatures from each data source were chosen. Altogether we generated 489 gene signatures corresponding to 64 cell types spanning multiple adaptive and innate immunity cells, hematopoietic progenitors, epithelial cells and extracellular matrix cells (Supplementary Table 2). Observing the scores in the 97 test primary cell types samples affirmed their ability to identify the corresponding cell type compared to other cell types across data sources (Supplementary Figure 2). We defined the raw enrichment score per cell type to be the average ssGSEA scores from all the cell type’s corresponding signatures.

### Spillover compensation between closely related cell types

Our primary objective is to accurately identify enrichments of cell types in mixtures. To imitate such admixtures, we performed an array of simulations of gene expression combinations of cell types to assess the accuracy and sensitivity of our gene signatures. We generated such *in silico* expression profiles using different data sources; using different sets of cell types in mixtures; and by choosing randomly one sample per cell type from all available samples in the data source. The simulations revealed that our raw scores reliably predict even small changes in the proportions of cell types, distinguish between most cell types, and are reliable in different transcriptomic analysis platforms (Supplementary Figure 3). However, the simulations also revealed that raw scores of RNA-seq samples are not linearly associated with the abundance and that they do not allow comparisons across cell types (Supplementary Figure 4). Thus, using the training samples we generated synthetic expression profiles by mixing the cell type of interest with another non-related cell types. We then fit a formula that transforms the raw scores to cell-type abundances. We found that the transformed scores showed resemblance to the known fractions of the cell types in simulations, thus allowing to compare scores across cell types, and not just across samples (Supplementary Figure 5).

The simulations also revealed another limitation of the raw scores: closely related cell-types tend to have correlating scores (Supplementary Figure 5). That is, scores may show enrichment for a cell type due to a ‘spillover effect’ between closely related cell types. This problem mimics the spillover problem in flow-cytometry, in which fluorescent signals correlate with each other due to spectrum overlaps. Inspired by the compensations method used in flow-cytometry studies [24], we leveraged our simulations to generate a spillover matrix that allows correcting for correlations between cell types. To better compensate for low abundances in mixtures we created a simulated dataset where each sample contains 25% of the cell type of interest and the rest from a non-related cell type and produced a spillover matrix, a representation of the dependencies of scores between different cell types.

Applying the spillover correction procedure on the pure cell types (Figure 2A) and simulated expression profiles (Figure 2B-C and Supplementary Figures 5-6) showed that this method was able to successfully reduce associations between closely related cell types. For example, we generated simulated mixtures using an independent data source of multiple cell types that was not part of the development of the method (GSE60424) [25], and used our method to infer the underlying abundances. We observed decent performance in recapitulating the cell types distributions (on the diagonal). However, before correcting for spillovers, there were wrong associations between CD4+ and CD8+ T-cells, as well as between Monocytes and Neutrophils. The spillover correction was able to reduce those associations significantly without harming the correlations on the diagonal (Figure 2B). In addition, we generated simulated mixtures using the training samples (Supplementary Figure 5) and the testing samples (Supplementary Figure 6). In the 18 simulated mixtures using the testing samples, we observed an overall average decrease of 17.1% in significant correlations off the diagonal (Figure 2C and Supplementary Figure 5). Unexpectedly, following the spillover compensation we observed slightly improved associations on the diagonal between the scores and the underlying abundances (1.4% average improvement). This pipeline of generating adjusted cell type enrichment scores from gene expression profiles, which we named *xCell*, is available as an R package and a simple web tool: http://xCell.ucsf.edu/

**Figure 2.**
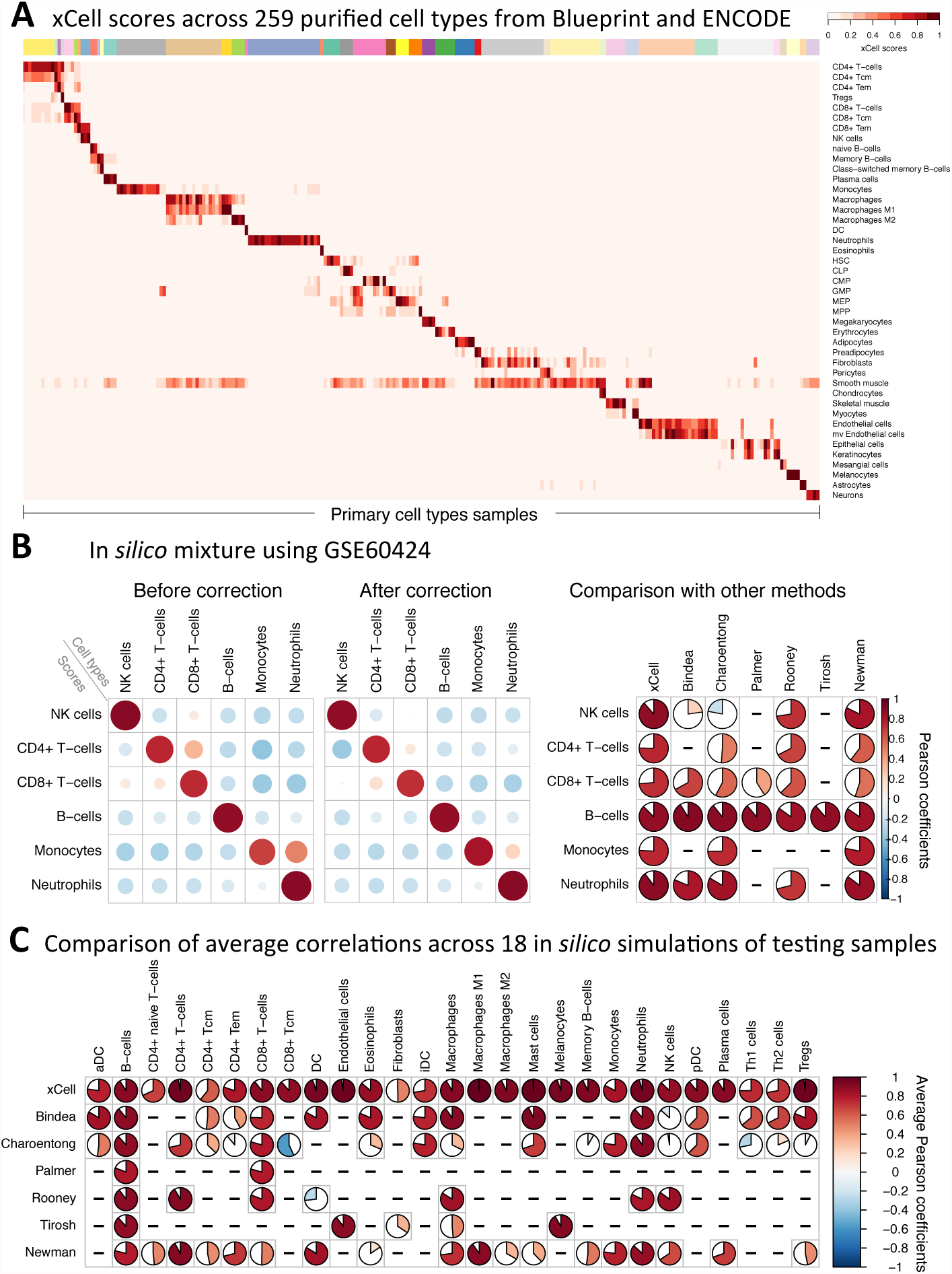
Evaluation of the performance of xCell using simulated mixtures. **A.** An overview of adjusted scores for of 43 cell type in 259 purified cell types samples from the Blueprint and ENCODE data sources (other data sources are in supplementary figure 4). Most signatures clearly distinguish the corresponding cell type from all other cell types. **B.** A simulation analysis using GSE60424 as the data source, which was not part of the development of xCell. This data source contains 114 RNA-seq samples from 6 major immune cell types. **Left:** Pearson correlation coefficients using our method before spillover adjustment and after the adjustment. Dependencies between CD4+ T-cells, CD8+ T-cells and NK cells were greatly reduced; spillover from monocytes to neutrophils was also removed. **Right:** Comparison of the correlation coefficients across the different methods. The first column corresponds to xCell’s inferences in predicting the underlying abundances of the cell types in the simulations (both color and pie chart correspond to average Pearson coefficients). Bindea, Charoentong, Palmer, Rooney and Tirosh represent sets of signatures for cell types from the corresponding manuscripts. Newman is the inferences produced using CIBERSORT on the simulations. xCell outperforms other methods in 17 of 18 comparisons. **C.** Comparison of the correlation coefficients across the different methods based on 18 simulations generated using the left-out testing samples. Here rows correspond to methods and columns show the average Pearson coefficient for the corresponding cell type across the simulations. Independent simulations are available in supplementary figure 6. xCell outperforms other methods in 64 of 67 comparisons.

We next compared the *xCell* scores ability to infer the underlying cell types enrichments in simulated mixtures with a set of 53 previously published signatures corresponding to 26 cell types [6,12,26,27] (Supplementary Table 3). Our analyses showed that *xCell* outperformed the previously published signatures in recapitulating the underlying abundances, in mixtures generated using the training samples (Supplementary Figure 5) and the testing samples (Supplementary Figure 6) and an independent data source (GSE60424) (Figure 2B), in the vast majority of the comparable cell types (51 of 53 comparisons of mixtures generated using trainings samples, 46 of 49 using testing samples and 17 of 18 using GSE60424) (Figure 2C). *xCell* showed overall better performance in all data sources used, proving its versatility across platforms. Importantly, our compensation technique was able to completely remove associations between cell types, while previously published signatures showed considerate dependencies between closely related cell types, such as between CD8+ T-cells and NK cells (Supplementary Figure 7).

In addition, we also compared *xCell’s* performance on test mixtures with CIBERSORT, a prominent deconvolution-based method [7]. Unlike signature-based methods, which output independent enrichment scores per cell type, the output from deconvolution-based methods is the inferred proportions of the cell types in the mixture. Similar to the performance compared to signatures, *xCell* also outperformed CIBERSORT enumerations in all comparable cell types, across all data sources (Figure 2B-C and Supplementary Figure 5-6).

### Validation of enrichment scores with cytometry immunoprofilings

In addition to the simulated mixtures analysis, we compared our estimates for cell types enrichments from gene expression profiles with mass spectrometry (CyTOF) immunophenotyping. We utilized independent publicly-available studies, in which a total of 165 individuals were studied for both gene expression from whole blood and FACS across 18 cell subsets from peripheral blood mononuclear cells (PBMC) (available in ImmPort SDY311 and SDY420) [28]. We calculated *xCell* scores for each of the signatures using the study’s expression profiles and correlated the scores with the FACS fractions of the cell subsets. Of the 14 cell types with at least 1% abundance, *xCell* was able to significantly recover 10 and 12 cell subsets in SDY311 and SDY420 respectively (Pearson correlation between calculated and actual cell counts *p-value* < 0.05) (Figure 3). Comparing the performance of *xCell* to previously published signatures and CIBERSORT revealed that no other method was able to recover cell types that our method was not able to recover in both data sets (Figure 3). In general, previous methods were able to recover signal only from major cell types, including B-cells, CD4+ and CD8+ T-cells, and monocytes, suggesting that their performance were not reliable in more specialized cell subsets. While our method also struggled in these cell subsets, it still showed significant correlations with most of the cell subsets, including effector memory CD8+ T-cells, naïve CD4+ T-cells and naïve B-cells. In addition, *xCell* was more reliable in CD4+ T-cells and monocytes and equally reliable in B-cells (Figure 3). In CD8+ T-cells *xCell* was outperformed by methods depending on solely on CD8A expression, which may not serve as a reliable biomarker in cancer settings (Supplementary Figure 7).

**Figure 3.**
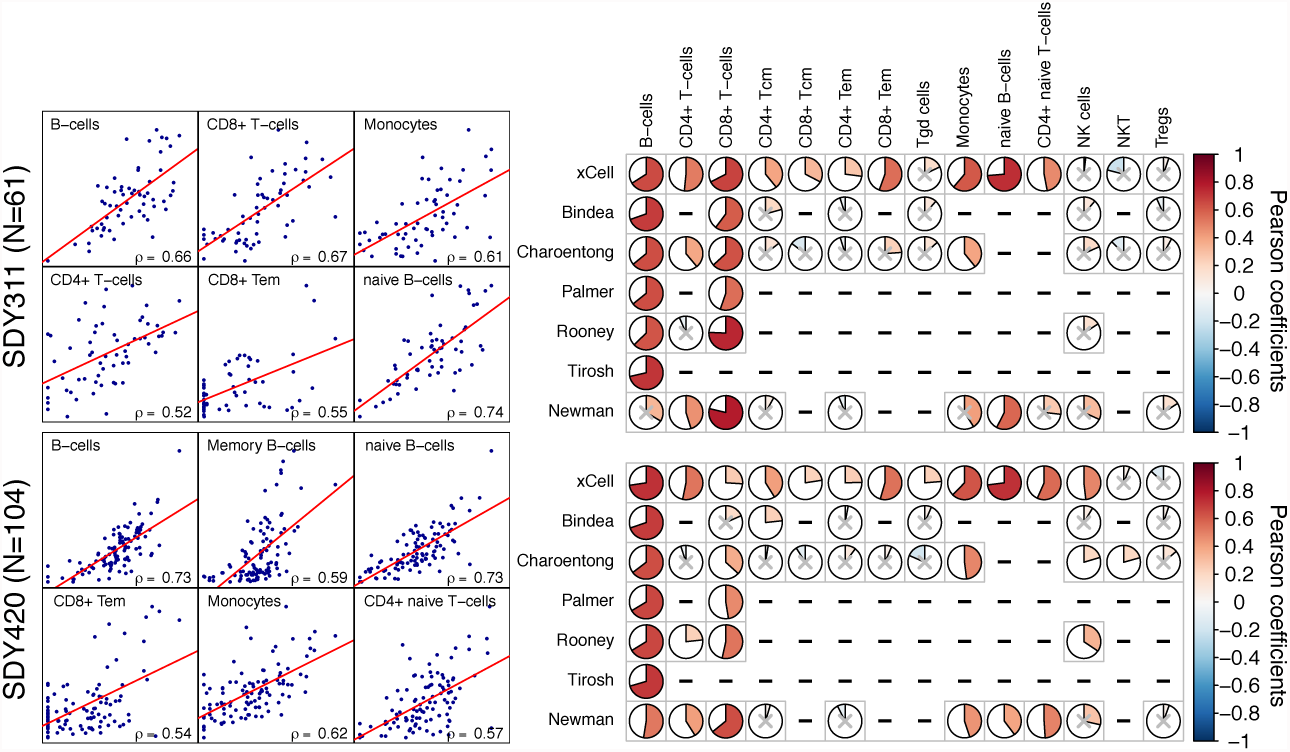
Compare digital dissection methods with flow cytometry counts. **Right:** Scatter plots of CyTOF fractions in PBMC vs. cell type scores from whole blood of 61 samples SDY311 (top) and 104 samples from SDY420 (bottom). Only top correlating cell types in each study is shown. **Left:** Correlation coefficients produced by our method compared to other methods. Only cell types with abundance of at least 1% on average, as measured by CyTOF, are shown. Non-significant correlations (p-value < 0.05) are marked with a gray ‘x’.

Despite the generally improved abilities of *xCell* to estimate cell populations, we do note that in some cases the correlations we observed were relatively low, emphasizing the difficulty of estimating cell subsets in mixed samples, and the need for cautious examination and further validation of findings.

### Cell types enrichments in tumor samples

We next applied our methodology on 9,947 primary tumor samples across thirty-seven cancer types from the TCGA and TARGET projects [29] (Supplementary Figure 9). Average scores of cell types in each cancer type affirmed prior knowledge of expected enriched cell types, validating the power of our method for identifying the cell type of origin of cancer types. For example, epithelial cells were enriched in carcinomas, keratinocytes in squamous cell carcinomas, mesangial cells in kidney cancers, chondrocytes in sarcoma, neurons in brain tumors, hepatocytes in hepatocellular carcinoma, melanocytes in melanomas, B-cells in B-cells lymphoma, T-cells in thymoma, myeloid cells in AML and lymphocytes in ALL (Figure 4A). While these results are expected, it is reassuring that *xCell* can be applied to and studied in human cancers.

**Figure 4.**
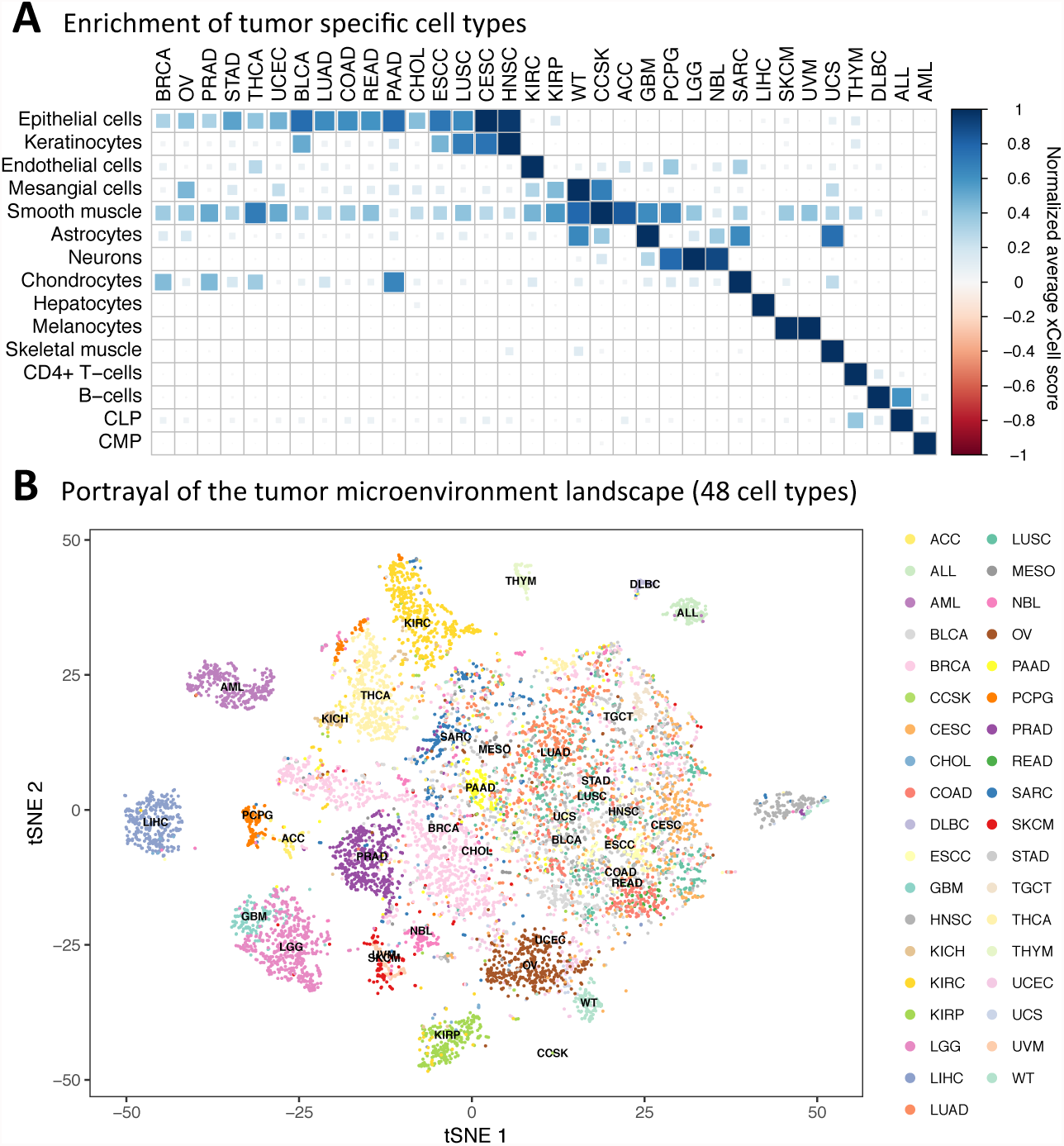
Cell types enrichment analysis in tumors. **A.** Average scores of 9 cell types across 24 cancer types from TCGA. Scores were normalized across rows. Signatures were chosen such that they are the cell of origin of cancer types or the most significant signature of the cancer type compared to all others. **B.** t-SNE plot of 8,875 primary cancer samples from TCGA colored by cancer type. The t-SNE plot was generated using the enrichment scores of 48 non-epithelial, non-stem cells and non-cell types specific scores. Many of the cancer types create distinct cluster, emphasizing the important role of the tumor microenvironment in characterizing tumors.

Most of the cell types we infer are part of the complex cellular heterogeneity of the tumor microenvironment. We hypothesized that an additive combination of all cell types’ scores would be negatively correlated with tumor purity. Thus, we generated a microenvironment score as the sum of all immune and stroma cell types. We then correlated this microenvironment score with our previously generated purity estimations, which are based on copy number variations, gene expression, DNA methylation and H&E slides [30]. Our analysis showed highly significant negative correlations in all cancer types, suggesting this score as a novel measurement for tumor microenvironment abundance (Supplementary Figure 10).

Finally, to provide insights into the potential of *xCell* to portray the tumor microenvironment, we plot all tumor samples based on their cell types scores. Using different sets of cell types inferences, we applied the t-Distributed Stochastic Neighbor Embedding (t-SNE) dimensionality reduction technique [31] (Supplementary Figure 11). Interestingly, the analysis revealed that unique microenvironment compositions characterize different cancer indications. For example, prostate cancers form a unique cluster based on their immune cell types composition, while head and neck tumors are distinguished by their stroma composition. Remarkably, only when performing the analysis with all immune and stroma cell types, clear clusters formed for distinguishing between most of the cancer types (Figure 4B), demonstrating the unique composition of the tumor microenvironment, which differs between cancer types. This notion emphasizes the importance of portraying the full cellular heterogeneity of the tumor microenvironment for the study of cancer. To this end, we calculated the enrichment scores for 64 cell types across the TCGA spectrum, and provide this data with the hope that it will serve the research community as a resource to further explore novel associations of cell types enrichment in human tumors (Supplementary Table 4).

## Discussion

Recently, many studies have shown different methodologies for digital dissection of cancer samples [3,6,9–13]. These studies have suggested novel insights in cancer research and related to therapy efficacy. However, it is important to remember that the methods that have been applied for portraying the tumor microenvironment have only retained limited validation, and it is unclear how reliable their estimations are. In this study, we took a step back and focused on generating cell type gene scores that could reliably estimate enrichments of cell types. Our method, which is gene-signature based, is more reliable due to its reliance on a group of signatures for each cell type, learned from multiple data sources, which increases the ability to detect the signal from the noise. Our method also integrates a novel approach to remove dependencies between cell types, which allow better reliability in studying closely related cell types.

To develop *xCell*, we collected the most comprehensive resource to date of primary cell types, spanning the largest set of human cell types. We then performed an extensive validation of the predicted cell types inferences in mixed samples. Our method for choosing a set of signatures that are reliable across several data sources has proven to be beneficial, as our scores robustly outperformed all available methods in predicting the abundance of cell types in *in silico* mixtures and blood samples. Based on our evaluation, *xCell* provides the most accurate and sensitive way to identify enrichments of many cell types in an admixture, allowing the detection of subtle differences in the enrichments of a particular cell type in the tumor microenvironment with high confidence.

It is important to note that *xCell*, as all other methods, performed significantly better in simulated mixtures than in real mixtures. Here, there are several technical reasons for this discrepancy. First, the cytometry analyses were performed on PBMCs, while the gene expression profiles were generated from the whole blood. Second, not all genes required by *xCell* were present, in fact in SDY420 only 54.5% of the genes required by *xCell* were available. However, other explanations for the lower success to infer abundances in real samples are warranting – it may well be possible that the expression patterns of marker genes in mixtures is different than in purified cells. Recent technologies such as single-cell RNA-sequencing may be able to clarify how much this may perturb the analyses.

We chose to apply a gene signature enrichment approach over deconvolution methods because of several advantages that the former provides. First, gene signatures are rank-based, and therefore are suitable for cross-platform transcriptome measurements. We showed here that our scores reliably predict enrichments in different RNA-seq techniques and different microarrays platforms. They are agnostic to normalization methods or concerns related to batch effects, making them robust to both technical and biological noise. Second, there is no decline in performance with increasing number of cell types. The tumor microenvironment is a rich milieu of cell types, and our analyses show enrichments of many mesenchymal derived cells in the tumors. A partial portrayal of the tumor microenvironment may result in misleading findings. Finally, gene signatures are simple and can easily be adjusted.

The main disadvantage of gene signatures is their difficulty to discriminate closely related cell types, though it is not clear how well other methods can distinguish such cell types as well [10]. Our method takes this into account and uses a novel technique, inspired by flow cytometry analyses, to remove such dependencies between closely related cell types. It is important to note that until this step the cell types scores are independent of each other, and a deflection of genes of one cell type will not harm all other cell types. However, the ‘spillover correction’ adjustment removes this strict independence between cell types inferences as in deconvolution methods. Yet, the compensation is very limited, and between most cell types there is no compensation at all, thus, most of the inferences are still independent.

Despite the utility of our signatures for characterizing the tumor microenvironment, several issues require further investigation. While our signatures outperformed previous methods, it is important to note that our correlations were still far from perfect with direct measurements. More expression data from pure cell types, especially cell types with limited samples, and more expression data coupled with cytometry counts from various tissue types will allow defining signatures more precisely and in turn, allow better reliability. Meanwhile, it is necessary to refer to inferences made by our method or other methods with a grain of salt. Discoveries made using digital dissection methods must be rigorously validated using other technologies to avoid hasty conclusions.

Another limitation of our method is that the inferences are strictly enrichment scores, and cannot be interpreted as proportions. This is due to the inability to translate the minimal and maximal scores produced by ssGSEA to clear proportions. Thus, while our method attempts to calibrate the scores to resemble proportions, this cannot be reliably used as such. This limitation also does not allow to provide statistical significance for the inferences, by calculating an empirical p-value as suggested by Newman et al. [7].

In summary, tissue dissection methods are an emerging tool for large-scale characterization of the tumor cellular heterogeneity. These approaches do not rely on tissue dissociation, opposed to single-cell techniques, and therefore provide an effective tool for dissecting solid tumors. The immense availability of public gene expression profiles allows these methods to be efficiently performed on hundreds of historical cohorts spanning thousands of patients, and to associate them with clinical outcomes. Here we presented the most comprehensive collection of gene expression enrichment scores for cell types. Our methodology for generating cell type enrichment scores and adjusting them to cell types proportions allowed us to create a powerful tool that is the most reliable and robust tool currently available for identifying cell types across data sources. We provide a simple web tool to the community and hope that further studies will utilize it for the discovery of novel predictive and prognostic biomarkers, and new therapeutic targets: http://xCell.ucsf.edu/.

## Methods

### Data sources

#### Signatures data sources

RNA-seq and cap analysis gene expression (CAGE) normalized FPKM were downloaded from the FANTOM5, ENCODE and Blueprint data portals. Raw Affymetrix microarray CEL files were downloaded from the Gene Expression Omnibus (GEO), accessions: GSE22886 (IRIS), GSE24759 (Novershtern) and GSE49910 (HPCA), and analyzed using the Robust Multi-array Average (RMA) procedure on probe-level data using Matlab functions. The analysis was performed using custom CDF files downloaded from Brainarray [32]. All samples were manually annotated to 64 cell types (Supplementary Table 1).

#### Other expression data sources

RNA-seq normalized counts were downloaded from the gene expression omnibus (GEO) accession GSE60424. Illumina HumanHT-12 V4.0 Beadchip data of PBMC samples and the accompanying CyTOF data were downloaded from ImmPort accession SDY311, and quantile normalized using Matlab functions. Similarly, Agilent Whole Human Genome 4 × 44 K slides data of PBMC samples and the accompanying CyTOF data were downloaded from ImmPort accession SDY420 [33], and quantile normalized using Matlab functions. Multiple probes per gene were collapsed using averages. RNA-seq data of Cancer Cell Line Encyclopedia (CCLE) [22] was obtained using the PharmacoGx R package [34]. RSEM levels 9,947 primary tumor samples from TCGA and TARGET were downloaded from https://toil.xenahubs.net. Published signatures were collected from their corresponding papers [6,12,26,27]. (Supplementary Table 3).

### *In silico* simulated mixtures

We generated several types of simulated mixtures, but all are based on the same pipeline:

1) Given a data source of pure cell types, choose *n* cell types available in the data and choose a random fraction for each cell type (the fractions sum to 1). We denote this vector of fraction *f*.
2) Generate an expression matrix of pure cell types, *M*, with *n* columns. The generation of the expression matrix varied between the experiments we performed: a) Synthetic mixtures for learning the power coefficient and spillover matrix were generated using the median expression profile of each cell type, creating a homogenous and noiseless mixture. b) Training mixtures were generated by randomly choosing one of the multiple available samples for each of the cell types chosen to be included in the mixture. This random selection introduces significant noise to the mixture, and between mixtures in the mixture set, which reflect the variation we observe between real datasets. c) Test mixtures, where only one sample per cell type was available, were generated by adding a random noise for each gene of up to 20% of the expression level. Cell types included in a mixture were chosen randomly, by avoiding cell types that cannot be distinguished (e.g. CD4+ T-cells and CD4+ memory T-cells).
3) To generate a simulated expression profile we use the formula: *M*×*f*, which returns one simulated mixed gene expression profile based on additive expression of the expression profiles of the cell types. This process is then repeated 500 times with different *f* and different *M* (as explained in 2b and 2c, *M* is recreated for each simulation by adding random noise (in b) or choosing a random sample), generating distinct mixtures using same set of cell types.

### The xCell development pipeline

A workflow of the xCell development pipeline can be found in Supplementary Figure 1, and is described in details below.

#### Filtering cancer genes

In a previous study [ 16] we calculated using CCLE the number of cell lines that are over-expressing each gene (2-fold more than the peak of expression distribution). For generating the signatures we only use genes that have an overexpression rate of less than 5% (less than 32 cell lines of the 634 carcinoma cell lines). We use this stringent threshold to eliminate genes that tend to be overexpressed in tumors, regardless of the cellular composition. Of 18,988 genes analyzed, 9,506 genes were identified as not overexpressing in tumors. For signatures of cell types that may be the cell of origin of solid tumors, including epithelial cells, sebocytes, keratinocytes, hepatocytes, melanocytes, astrocytes and neurons we used all genes.

#### Generating gene signatures

Expression profiles were reduced to 10,808 genes that are shared across all 6 data sources. Gene expression was converted to log scale by adding 3 and then log_2_ conversion. In each group of samples corresponding to a cell type we calculated 10^th^, 25^th^, 33.3^th^, 50^th^ percentiles of low expression (*Q1_q_*), and 90^th^, 75^th^, 66.6^th^, 50^th^ quantiles of high expression (*Q2*_1*-q*_). For cell type *A* we calculated the difference for each gene between *Q1*_q_(A) and max(*Q2*_1-q_(all other cell types)). We repeated this also for second and third largest*Q2*_1-q_(all other cell types). The signature of cell type *A* consists of all genes that pass a threshold. We used different thresholds here: 0, 0.1, log2(1.5), log(2), 3,4 and 5. We repeated this procedure to each of the 6 data sources independently. Only gene sets of at least 8 genes and no more than 200 genes were reserved. This scheme yielded 6,573 gene signatures corresponding to 64 cell types. We calculated single-sample gene set enrichment analysis (ssGSEA) for each of those signatures to score each sample in each of the data sources using the GSVA R package [35].

#### Choosing the “best” signature

For each signature we computed the t-statistic between the scores of the corresponding cell type compared to all other samples, omitting samples from parental or descendant cell types (for CD4+ naïve T-cells the general CD4+ T-cells are not used in the calculations). The procedure was performed in each data source where the corresponding cell type was available, except the data source from which the signature was learned. Thus, a signature is only chosen if it is reliable in a data source were it was not trained upon. If the cell type is available in only one data source, the signature was tested in that data source. From each data source the top 3 signatures were chosen. All together we chose 489 signatures corresponding to 64 cell types (across the 6 data sources we have 163 cell types) (Supplementary Table 2). The raw score for a cell type is the average of all corresponding signatures, after shifting scores of each signature to have a minimal score of 0 across all samples.

#### Learning parameters for raw score transformation

For each cell type we created a synthetic mixture using the median expression profile of the cell type (cell *X*) and an additional ‘control’ cell type. For ‘control’ in sequencing-based data sources we used multipotent progenitor (MPP) cell samples or endothelial cell samples, because both are found in all the sequencing-based data sets. In microarray-based data sources we used erythrocytes and monocytes. We generated such mixtures using increasing levels of the corresponding cell type (0.8% of the cell *X* and 99.2% control, 1.6% cell *X* and 98.4% control, etc.). We noticed two problems with the raw scores: ssGSEA scores have different distributions between different signatures, thus a score from signatures cannot be compared with a score from another signature. In addition, in sequencing-based data, the association between the underlying levels of the cell type was not linearly associated with the score. We thus designed a transformation pipeline for the scores (which is applied to both sequencing and microarray-based datasets separately) – for each cell type, using the synthetic mixtures we first shifted the scores to 0 using the minimal score (which corresponded to mixtures containing 0.8% of the cell type) and divided by 5000. We then fit a power function to the scores ranging corresponding to abundances of 0.8% to 25.6%. We used this range because we are mostly interested in identifying cell types with low abundance, and above that the function exponential increase may interfere in a precise fitting. The power coefficient is then averaged across the data sources were the cell type is available (we denote this vector as P). After adjusting the score using the learned power coefficient, we fit a linear curve, and use the learned slope as a calibration parameter for the adjusted scores (denoted as *V*1).

#### Learning the spillover compensation reference matrix

Another limitation that was observed in the mixtures is the dependencies between closely related cell types: scores that predict enrichment of one cell type also predict enrichment of another cell type, which might not even be in the mixture. To overcome this problem we created a reference matrix of ‘spillovers’ between cell types. Below we focus on the generation of the sequencing-based spillover matrix, however equivalent process was performed to generate the microarray-based spillover matrix. We first generated a synthetic mixture set, where each mixture contains 25% of each of the cell types (median expression) and 75% of a ‘control’ cell type, as in the previous section. We then calculated raw cell types scores and transformed them using the learned coefficients as explained above. We combined all sequencing-based data sources together by using the average scores, and completed the matrix to be 64×64 by adding columns from cell types that are not present in any of the sequencing-based data using the microarray reference matrix. We then normalized each row of cell types scores by dividing it by the diagonal (denoted as *K* – spillover matrix, rows are cell types scores and columns are cell type samples). The diagonal, before the normalization, is also used for calibration (denoted as *V*2) The ‘spillover’ between a cell type score (*x*) and another cell type (*y*) is the ratio between *x* and *y*. Finally, we clean the spillover matrix to not compensate between parent and descendent cell types by compensating parent cell types only with other parent cell types (CD4+ T-cells are compensated against CD8+ T-cells, but not CD8+ Tem), and compensating child cell types only compared to other child cell types from the same parent and all other parents, but not child cell types from other parents. Some of the compensations where too strong, removing correlations between cell types and their corresponding signatures, thus we limit the compensation levels, off the diagonal, to 0.5. The spillover matrix, power and calibration coefficients are available in Supplementary Table 5.

#### Calculating scores for a mixture

Input: Gene expression data set normalized to gene length (such as FPKM or TPM), rows are genes and columns are samples (N – number of samples). Duplicate gene names are combined together. *xCell* uses a set of 10,808 genes for the scoring. It is recommended to use data sets that contain at least the majority of these genes. Missing values in a sample are treated as missing genes. It is also recommended to use as many samples as possible, with highly expected variation in cell types fractions (the xCell web tool requires intersection of at least 5000 genes). (1) Calculating ssGSEA for each of the 489 gene signatures. (2) Averaging scores of all signatures corresponding to a cell type. The result is a matrix (*A*) with 64 rows and N columns. (3) Each element in the scores matrix (*A_ij_*) is transformed using the following formula:

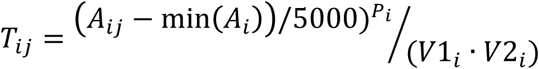

The output is matrix *T* of transformed scores. Different P, V1 and V2 are used for sequencing-based and microarray-based datasets. (4) Spillover compensation is then performed for each row using linear least squares that minimizes the following (as performed in flow cytometry analyses and explained in Bagwell et al. [24]):

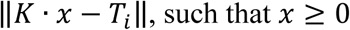

All *x*’s are then combined to create the final xCell scores. The compensation may result in deteriorating real associations, thus we provide a scaling parameter (alpha), to multiply all off-diagonal cells in matrix K. In all experiments in the manuscript we used alpha=0.5. Different K matrices are used for sequencing-based and array-based data.

### Cytometry analyses

Gene expression and cytometry data were downloaded from ImmPort (SDY311 and SDY420). The gene expression data was quantile normalized using Matlab functions, and multiple probes per gene were collapsed using averages. The cytometry data counts were divided by the viable/singlets counts. In the SDY311 dataset 10 patients had two replicates of expression profiles, and those were averaged. Two outlier samples in the cytometry data set were removed from further analyses (SUB134240, SUB134283).

### Other tools

The CIBERSORT web tool was used for inferring proportions using the expression profile (http://cibersort.stanford.edu). CIBERSORT results of activated and resting cell types were combined, B-cell and CD4+ T-cells percentages are the combination of all their subtypes. t-SNE plots were produced using the Rtsne R package. Purity measurements were obtained from our previous publication [30]. Correlation plots were generated using the corrplot R package.

### Availability of Data and Materials

The xCell R package for generating the cell type scores, and R scripts for the development of xCell are available at https://github.com/dviraran/xCell, and deposited to Zenodo (assigned DOI: 10.5281/zenodo.800745).

## Supplementary Information

**Supplementary Figure 1. The xCell development pipeline.** A schematic illustration of the steps performed for the development of xCell.

**Supplementary Figure 2. Raw scores of the multiple signatures in the test samples.** Box plots of the raw scores of all signatures corresponding to a cell type presented on the 97 testing cell types. xCell uses the average of the raw scores for downstream analyses.

**Supplementary Figure 3. Simulated mixtures of pure cell types inferred by raw xCell scores.** Scatter plots of inferred vs. simulated abundance in 10 mixtures sets.

**Supplementary Figure 4. Transformation procedure of raw scores to linear scales.** The associations between raw scores and simulated abundances before and after transformation.

**Supplementary Figure 5. Cell types inferences in gene expression simulations using training samples.** Simulations analyses using training samples, showing the correlation between cell types based on xCell inferences compared to other methods.

**Supplementary Figure 6. Cell types inferences in gene expression simulations using testing samples.** Simulations analyses using the left-out testing samples, showing the correlation between cell types based on xCell inferences compared to other methods.

**Supplementary Figure 7. Dependencies between CD8+ T-cells and NK cells**. An example of dependency between cell types in published signatures.

**Supplementary Figure 8. CD8+ T-cells scores vs. CD8A expression in cancer cell lines.** An example of the problem of using a single gene for predicting cell type abundance.

**Supplementary Figure 9. xCell scores in 24 TCGA cancer types.** Box plots of cell types scores across 24 tumors types from TCGA.

**Supplementary Figure 10. Purity estimations using xCell scores.** Comparing a tumor microenvironment derived by xCell to the CPE purity estimation.

**Supplementary Figure 11. t-SNE plots based on cell types scores.**

**Supplementary Table 1. Summary table of primary cell types used in this study**. Number of samples annotated for each of the 64 cell types across the 6 data sources.

**Supplementary Table 2. 489 cell type gene signatures.**

**Supplementary Table 3. 53 previously published cell type gene signatures.**

**Supplementary Table 4. xCell scores in 9,264 samples from TCGA.**

**Supplementary Table 5. Spillover matrix and calibrating coefficients.**

## Acknowledgments

We thank Marina Sirota and Thomas Peterson for helpful discussions. This work was supported by the Gruss Lipper Postdoctoral Fellowship to D.A., and the National Cancer Institute (U24 CA195858) and the National Institute of Allergy and Infectious Diseases (Bioinformatics Support Contract HHSN272201200028C) to A.J.B. The content is solely the responsibility of the authors and does not necessarily represent the official views of the National Institutes of Health.

## References

1. Galon J, Costes A, Sanchez-Cabo F, Kirilovsky A, Mlecnik B, Lagorce-Pagès C, et al. Type, density, and location of immune cells within human colorectal tumors predict clinical outcome. Sci. (New York, NY) [Internet]. 2006;313:1960–4. Available from: papers3://publication/doi/10.1126/science.1129139

2. Hanahan D, Coussens LM. Accessories to the Crime: Functions of Cells Recruited to the Tumor Microenvironment. Cancer Cell. 2012. p. 309–22.

3. Gentles AJ, Newman AM, Liu CL, Bratman S V, Feng W, Kim D, et al. The prognostic landscape of genes and infiltrating immune cells across human cancers. Nat. Med. [Internet]. 2015;21:938–45. Available from: http://www.ncbi.nlm.nih.gov/pubmed/26193342

4. Abbas AR, Wolslegel K, Seshasayee D, Modrusan Z, Clark HF. Deconvolution of blood microarray data identifies cellular activation patterns in systemic lupus erythematosus. PLoS One. 2009;4.

5. Shen-Orr SS, Gaujoux R. Computational deconvolution: extracting cell type-specific information from heterogeneous samples. Curr. Opin. Immunol. [Internet]. 2013 [cited 2015 Mar 24];25:571–8. Available from: http://www.sciencedirect.com/science/article/pii/S0952791513001507

6. Rooney MS, Shukla SA, Wu CJ, Getz G, Hacohen N. Molecular and genetic properties of tumors associated with local immune cytolytic activity. Cell. 2015;160:48–61.

7. Newman AM, Liu CL, Green MR, Gentles AJ, Feng W, Xu Y, et al. Robust enumeration of cell subsets from tissue expression profiles. Nat. Methods [Internet]. 2015 [cited 2015 Mar 30];12:453–7. Available from: http://www.ncbi.nlm.nih.gov/pubmed/25822800

8. Newman AM, Alizadeh AA. High-throughput genomic profiling of tumor-infiltrating leukocytes. Curr. Opin. Immunol. 2016;41:77–84.

9. Angelova M, Charoentong P, Hackl H, Fischer ML, Snajder R, Krogsdam AM, et al. Characterization of the immunophenotypes and antigenomes of colorectal cancers reveals distinct tumor escape mechanisms and novel targets for immunotherapy. Genome Biol. [Internet]. 2015;16:64. Available from: http://genomebiology.com/2015/16/1/64

10. Li B, Severson E, Pignon J-C, Zhao H, Li T, Novak J, et al. Comprehensive analyses of tumor immunity: implications for cancer immunotherapy. Genome Biol. 2016;17:14.

11. Iglesia MD, Parker JS, Hoadley KA, Serody JS, Perou CM, Vincent BG. Genomic Analysis of Immune Cell Infiltrates Across 11 Tumor Types. J. Natl. Cancer Inst. [Internet]. Oxford University Press; 2016 [cited 2016 Aug 10];108:djw144. Available from: http://jnci.oxfordjournals.org/lookup/doi/10.1093/jnci/djw144

12. Charoentong P, Finotello F, Angelova M, Mayer C, Efremova M, Rieder D, et al. Pan-cancer Immunogenomic Analyses Reveal Genotype-Immunophenotype Relationships and Predictors of Response to Checkpoint Blockade. Cell Rep. 2017;18:248–62.

13. Şenbabaoğlu Y, Gejman RS, Winer AG, Liu M, Van Allen EM, de Velasco G, et al. Tumor immune microenvironment characterization in clear cell renal cell carcinoma identifies prognostic and immunotherapeutically relevant messenger RNA signatures. Genome Biol. [Internet]. 2016 [cited 2017 Feb 1]; 17:231. Available from: http://genomebiology.biomedcentral.com/articles/10.1186/s13059-016-1092-z

14. Pattabiraman DR, Weinberg RA. Tackling the cancer stem cells - what challenges do they pose? Nat. Rev. Drug Discov. [Internet]. 2014;13:497–512. Available from: http://www.ncbi.nlm.nih.gov/pubmed/24981363%5Cnhttp://dx.doi.org/10.1038/nrd4253

15. Turley SJ, Cremasco V, Astarita JL. Immunological hallmarks of stromal cells in the tumour microenvironment. Nat. Rev. Immunol. [Internet]. 2015;15:669–82. Available from: http://www.ncbi.nlm.nih.gov/pubmed/26471778

16. Aran D, Lasry A, Zinger A, Biton M, Pikarsky E, Hellman A, et al. Widespread parainflammation in human cancer. Genome Biol. [Internet]. BioMed Central; 2016 [cited 2016 Jul 21];17:145. Available from: http://genomebiology.biomedcentral.com/articles/10.1186/s13059-016-0995-z

17. Lizio M, Harshbarger J, Shimoji H, Severin J, Kasukawa T, Sahin S, et al. Gateways to the FANTOM5 promoter level mammalian expression atlas. Genome Biol. [Internet]. 2015;16:22. Available from: http://genomebiology.com/2015/16/1/22

18. Consortium EP, Bernstein BE, Birney E, Dunham I, Green ED, Gunter C, et al. An integrated encyclopedia of DNA elements in the human genome. Nature [Internet]. 2012;489:57–74. Available from: http://www.nature.com/doifinder/10.1038/nature11247%5Cnpapers3://publication/doi/10.1038/nature11247

19. Abbas AR, Baldwin D, Ma Y, Ouyang W, Gurney A, Martin F, et al. Immune response in silico (IRIS): immune-specific genes identified from a compendium of microarray expression data. Genes Immun. [Internet]. 2005;6:319–31. Available from: http://www.ncbi.nlm.nih.gov/pubmed/15789058

20. Novershtern N, Subramanian A, Lawton LN, Mak RH, Haining WN, McConkey ME, et al. Densely interconnected transcriptional circuits control cell states in human hematopoiesis. Cell. 2011;144:296–309.

21. Mabbott NA, Baillie JK, Brown H, Freeman TC, Hume DA. An expression atlas of human primary cells: inference of gene function from coexpression networks. BMC Genomics [Internet]. 2013;14:632. Available from: http://www.pubmedcentral.nih.gov/articlerender.fcgi?artid=3849585&tool=pmcentrez&rendertype=abstract

22. Barretina J, Caponigro G, Stransky N, Venkatesan K, Margolin AA, Kim S, et al. The Cancer Cell Line Encyclopedia enables predictive modelling of anticancer drug sensitivity. Nature [Internet]. 2012;483:603–7. Available from: http://dx.doi.org/10.1038/nature11003

23. Barbie DA, Tamayo P, Boehm JS, Kim SY, Moody SE, Dunn IF, et al. Systematic RNA interference reveals that oncogenic KRAS-driven cancers require TBK1. Nature [Internet]. 2009;462:108–12. Available from: http://www.pubmedcentral.nih.gov/articlerender.fcgi?artid=2783335&tool=pmcentrez&rendertype=abstract

24. Bagwell CB, Adams EG. Fluorescence spectral overlap compensation for any number of flow cytometry parameters. Ann. N. Y. Acad. Sci. [Internet]. 1993 [cited 2017 Feb 5];677:167–84. Available from: http://www.ncbi.nlm.nih.gov/pubmed/8494206

25. Linsley PS, Speake C, Whalen E, Chaussabel D. Copy Number Loss of the Interferon Gene Cluster in Melanomas Is Linked to Reduced T Cell Infiltrate and Poor Patient Prognosis. Castro MG, editor. PLoS One [Internet]. 2014 [cited 2017 Feb 2];9:e109760. Available from: http://www.ncbi.nlm.nih.gov/pubmed/25314013

26. Bindea G, Mlecnik B, Tosolini M, Kirilovsky A, Waldner M, Obenauf AC, et al. Spatiotemporal dynamics of intratumoral immune cells reveal the immune landscape in human cancer. Immunity. 2013;39:782–95.

27. Tirosh I, Izar B, Prakadan SM, Wadsworth MH, Treacy D, Trombetta JJ, et al. Dissecting the multicellular ecosystem of metastatic melanoma by single-cell RNA-seq. Science (80-. ). [Internet]. 2016;352:189–96. Available from: http://science.sciencemag.org.gate2.inist.fr/content/352/6282/189.abstract

28. Bhattacharya S, Andorf S, Gomes L, Dunn P, Schaefer H, Pontius J, et al. ImmPort: Disseminating data to the public for the future of immunology. Immunol. Res. 2014;58:234–9.

29. Vivian J, Rao AA, Nothaft FA, Ketchum C, Armstrong J, Novak A, et al. Toil enables reproducible, open source, big biomedical data analyses. Nat. Biotechnol. 2017;

30. Aran D, Sirota M, Butte AJ. Systematic pan-cancer analysis of tumour purity. Nat. Commun. [Internet]. 2015;6:8971. Available from: http://www.nature.com/doifinder/10.1038/ncomms9971

31. van der Maaten L, Hinton GE. Visualizing high-dimensional data using t-SNE. J. Mach. Learn. Res. [Internet]. 2008;9:2579–605. Available from: http://www.ncbi.nlm.nih.gov/entrez/query.fcgi?db=pubmed&cmd=Retrieve&dopt=AbstractPlus&list_uids=7911431479148734548related:VOiAgwMNy20J

32. Dai M, Wang P, Boyd AD, Kostov G, Athey B, Jones EG, et al. Evolving gene/transcript definitions significantly alter the interpretation of GeneChip data. Nucleic Acids Res. 2005;33.

33. Whiting CC, Siebert J, Newman AM, Du H, Alizadeh AA, Goronzy J, et al. Large-Scale and Comprehensive Immune Profiling and Functional Analysis of Normal Human Aging. Unutmaz D, editor. PLoS One [Internet]. 2015 [cited 2017 Feb 6];10:e0133627. Available from: http://www.ncbi.nlm.nih.gov/pubmed/26197454

34. Smirnov P, Safikhani Z, El-Hachem N, Wang D, She A, Olsen C, et al. PharmacoGx: an R package for analysis of large pharmacogenomic datasets. Bioinformatics [Internet]. 2016 [cited 2017 Feb 6];32:1244–6. Available from: http://www.ncbi.nlm.nih.gov/pubmed/26656004

35. Hänzelmann S, Castelo R, Guinney J. GSVA: gene set variation analysis for microarray and RNA-seq data. BMC Bioinformatics [Internet]. 2013;14:7. Available from: http://www.pubmedcentral.nih.gov/articlerender.fcgi?artid=3618321&tool=pmcentrez&rendertype=abstract

